# A well-ordered nanoflow LC-MS/MS approach for proteome profiling using 200 cm long micro pillar array columns

**DOI:** 10.1101/472134

**Authors:** Jeff Op De Beeck, Jarne Pauwels, Natalie Van Landuyt, Paul Jacobs, Wim De Malsche, Gert Desmet, Andrea Argentini, An Staes, Lennart Martens, Francis Impens, Kris Gevaert

## Abstract

In bottom-up proteomics, capillaries up to 75 cm long with internal diameters of 50 to 100 µm packed with sub-2-µm C18-functionalized particles are routinely used in combination with high-resolution mass spectrometry. Unlike such conventional liquid chromatography (LC) columns, micro pillar array columns (µPAC™) are fabricated using micromachining technology, resulting in perfectly ordered chromatographic separation beds, leading to a minimized analyte dispersion while column permeability is increased by one order of magnitude. This allows using very long columns (up to 200 cm) at only a fraction of the pressure needed to operate packed bed columns. To validate µPAC™ column performances, different amounts of tryptic digests of HEK293T cell lysates were prepared and separated using a 200 cm µPAC™ column or a 40 cm long conventional column. Using an Orbitrap Elite instrument, on average 25% more proteins were identified with the µPAC™ column. Moreover, the rate at which the peak width increases with gradient time is much lower on the µPAC™ column. For a 10-hour long gradient, average peak widths below 0.5 min were observed, resulting in consistent identification of over 5,000 proteins. Combining long solvent gradients and this new type of LC column, substantial improvements in proteome coverage could be obtained. Finally, we demonstrated high reproducibility and durability of the µPAC™ column. Data are available via ProteomeXchange with identifiers PXD011547 and PXD013235.

---

Mass spectrometry (MS)-based proteomics has become an essential tool to provide biologists with accurate information about protein levels, posttranslational modifications and protein interaction partners^1^. With the ultimate goal to identify all proteoforms^2^ present in a biological sample, the practice of MS-based proteomic profiling has evolved enormously during the last decade^3^. Liquid chromatography coupled to tandem mass spectrometry (LC-MS/MS) still is the dominant technique for proteomic profiling and is continuously evolving to retrieve more data in shorter time^4^, while simultaneously decreasing operational complexity and increasing method robustness^5^. Whereas two decades ago, state-of-the-art MS systems were able to acquire up to one MS/MS scan per second, technological advances in MS instrumentation are nowadays capable of acquiring well over tens of MS/MS scans per second^6, 7^ The number of peptides and proteins that can be identified in one LC-MS/MS run clearly depends to a large extent on the scanning speed of the MS instrument, but equally depends on the quality of the chromatographic separation^8^. Especially now that MS instruments have become so fast, the main limiting factor appears to be chromatographic performance or resolving power^9^.

Chromatographic performance is commonly expressed as peak capacity (nC) which takes both the average peak width and the elution window into account. Peak capacity is a theoretical measure for the number of peaks that can be separated within a given timeframe and previous studies have pointed out a linear relation to the amount of peptides that can be identified^8, 10^. With documented values ranging from several hundred up to 1,500, peak capacity mainly depends on the properties of the LC column, but it is also affected by the conditions used to perform gradient elution chromatography such as column temperature, flow rate and gradient time^11–13^. When focusing on the column properties, an increase in peak capacity can be achieved by extending the column length or by decreasing the plate height (h). The plate height (or efficiency) of a column depends on the diameter of the particles that are used to pack the column and the packing quality^14^. Reducing the particle diameter will have a beneficial effect on chromatographic performance and will lead to higher peak capacities. However, reducing the particle diameter and extending the column length comes at the cost of increased backpressure. The pressure drop of a column is linearly related to its length and inversely proportional to the square of the particle diameter. As a consequence, current state-of-the art nano LC columns often require ultra-high performance liquid chromatography (UPLC) type of LC instruments which can accurately deliver flow rates of 0.05 to 1 µl/min at operating pressures up to 1,200 bar. Such columns are typically quite long, with lengths up to 50 cm and more, and packed with sub-2-µm C18 functionalized porous silica particles, producing peak capacity values between 600 and 800 for very long gradient separations (> 240 min). Peak capacities ranging from 1,000 to 1,500 have however been obtained when using longer columns or columns packed with even smaller particles, using custom built LC instrumentation capable of delivering operating pressures up to 1,360 bar^11, 15, 16^. Operating columns at pressures exceeding 1,000 bar is not straightforward as column fittings and LC instrument parts have to endure much more, hereby possibly compromising system robustness. A strategy to circumvent the increased backpressure that is observed by extending column lengths beyond 100 cm is the use of monolithic stationary phase support structures as an alternative to silica particles. Both silica and polymer-based monolithic columns have been developed and several of them are commercially available^17^. Due to the high permeability that is observed for monolithic columns, the use of columns with lengths exceeding 300 cm has been reported and such columns have been applied successfully to perform deep proteome profiling at operating pressures well below 400 bar^18, 19^.

Following up on the pioneering work of the Regnier lab^20, 21^, micro pillar array columns (µPAC™s) were introduced about two decades ago as a powerful alternative for classical packed bed columns and monoliths^22–26^. By using micro fabrication technology initially developed for the microelectronics industry, the stationary phase backbone of pillar array columns is precisely defined in high purity silicon wafers. The main advantage of this approach is that such columns are reproducibly manufactured with a high degree of uniformity and in a perfectly ordered manner, i.e. the stationary phase morphology (or bed of micro pillars) is uniform over the entire column width and length (Figure S-1, Supporting Information).

Due to the high degree of uniformity, peak dispersion originating from heterogeneous flow paths in the separation bed is eliminated (i.e. no Eddy dispersion), and therefore components remain much more concentrated during separation^27, 28^. Apart from an improved efficiency, pillar array columns have substantially lower flow resistances compared to packed bed columns. When packing spherical particles into a cylindrical column or capillary, the relative volume that is taken by the particles to that of the inter particle void space (called external porosity ε) will always be restricted to a single value of 0.4. This is the case for all packed bed columns, no matter what particle size or capillary diameter is used. The distance between the pillars can however be independently controlled from the pillar size, enabling the fabrication of columns over a range of external porosities. The higher the external porosity, the lower the flow resistance. Because of this low flow resistance, moderate LC pump pressures (below 350 bar) can be used to operate very long separation channels with lengths exceeding 100 cm. To fabricate very long columns on a silicon wafer with limited dimensions, the separation channel is folded in a serpentine manner. By interconnecting a total of 40 5-cm long separation channels with low dispersion flow distributor and turn structures, a column with a total length of 200 cm can be fabricated (Figure 1)^29, 30^.

**Figure 1.**
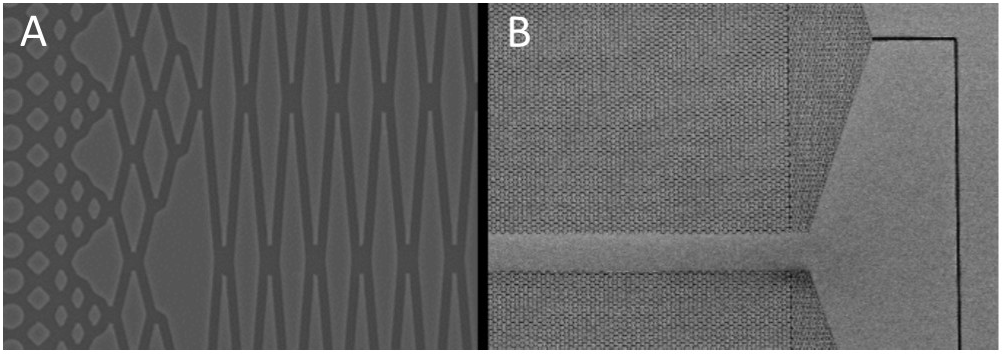
Scanning electron microscopy (SEM) images showing flow distributor structures that are used to interconnect parallel separation lanes of the micro pillar array column. A, Top view of the proprietary flow distribution design. B, Two of these structures are used to interconnect parallel separation lanes with a narrow 180° turn channel.

To investigate the benefits of the µPAC™ technology for RPLC-MS/MS based proteomics workflows, a series of benchmarking experiments were performed by injecting different amounts of tryptic protein digests of HEK293T cell lysates and separating these samples using an extensive range of solvent gradients. Employing an Orbitrap Elite mass spectrometer for MS/MS detection, we aimed at getting a clear assessment of both the chromatographic performance of this column type as well as its relation to the number of proteins and peptides that can be identified in a typical bottom-up proteomics experiment. For comparison, all experiments were repeated on a 40 cm long nano LC column packed in-house with sub 2 µm C18 functionalized porous silica particles which is considered to be among the top performing nano LC columns for the separation of complex peptide mixtures^3^. Finally, we showed high durability and reproducibility of µPAC™ columns when used for routine measurements on a Q Exactive HF instrument.

## Experimental Section

### Cell Culture

Human HEK293T cells were purchased from the ATCC (TIB-152, American Type Culture Collection, Manassas, VA, USA) and grown in Dulbecco’s modified eagle medium (DMEM) with Glutamax (Invitrogen, 31966-047) supplemented with 10% dialyzed foetal bovine serum (Gibco) and 1% Penicillin-streptomycin (Gibco, 5,000 U/mL). Cell pellets containing 5 million cells were harvested, washed with phosphate buffered saline (PBS) and frozen at −80°C until further use.

### Columns

The first analytical column was a classical packed bed column (40 cm long, 75 µm inner diameter) packed inhouse with ReproSil-Pur basic 1.9 µm silica particles (Dr. Maisch). Prior to packing the column, the fused silica capillary had been equipped with a laser pulled electrospray tip using a P-2000 Laser Based Micropipette Peller (Sutter Instruments). The second column was a 200 cm long micro pillar array column (µPAC™, PharmaFluidics) with C18-endcapped functionality. This column consists of 300 µm wide channels that are filled with 5 µm porous-shell pillars at an inter pillar distance of 2.5 µm. With a depth of 20 µm, this column has a cross section equivalent to an 85 µm inner diameter capillary column.

### Sample preparation, LC-MS/MS analysis and database searching

Different amounts of peptide material have been analyzed on both types columns, in duplicate and using different solvent gradients (see Figures). Full experimental details are available as Supporting Information.

### Data analysis

The comparison between the columns was primarily based on the overall number of both protein and peptide identifications, the difference in column pressure, evolution of peak width and overall sensitivity. The database search results were processed in Perseus (v1.6.1.3) after loading the proteinGroups file from MaxQuant^31^. For the shotgun samples derived from HEK293T cells, we first removed the reversed hits, potential contaminants and proteins that were only identified by site. The intensities were then log2 transformed and a Venn diagram was created, returning the number of identifications per run. Peak widths from several runs were extracted by the mOFF algorithm^32^ from each raw file and corresponding evidence.txt file. The parameters were set to 10 ppm peptide mass tolerance, 1 min retention time window for the extracted ion chromatogram (XIC) (default set to 3 min) and 0.1 min retention time window for the peak. Extracted peak widths at FWHM were converted to their corresponding value at 4σ using the relation 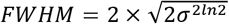 and the average peak width of all identified peptides was calculated. Peak capacity values were calculated using the equation *n*_*c*_ = (*T*_*G*_/*w*_4*σ*_) + 1 with n_C_ the peak capacity, T_G_ the solvent gradient time and w_4σ_ the average peak width at 4σ.

### Evaluation of the durability and reproducibility of the µPAC™ column

A commercial K562 cell digest supplemented with the 6 x 5 LC-MS/MS Peptide Reference Mix (QC4Life Reference Standard, Promega) was used as quality control (QC) to test the reproducibility of the µPAC™ column when used for routine measurements on a Q Exactive-type instrument over a 6-weeks period. Full experimental details are available as Supporting Information.

## Results and Discussion

In this study, we aimed at investigating the benefits that the µPAC™ column technology can offer for bottom-up proteomics. A concentration range of HEK293T tryptic digest samples (0.1-0.5-1-2-3 µg) was injected in duplicate and separated using solvent gradient times ranging from 30 to 570 min. By performing LC-MS/MS analysis, we intended to get a clear evaluation of the chromatographic performance and its effect on the number of peptide and protein identifications.

When comparing the 200 cm µPAC™ column to a classical packed bed column with a length of 40 cm, the first thing that stands out is the column pressure needed to operate this new type of column. Whereas classical packed bed nano LC columns packed with 1.9 µm porous particles easily generate pressure drops exceeding 400 bars, only 75 bar is needed to operate the µPAC™ column at a flow rate of 300 nl/min (Figure 2). This low back pressure can be attributed to the high permeability that is inherent to the pillar array column format and that is also observed for several types of monolithic columns^17–19^ and columns packed with porous-shell beads^33^.

**Figure 2.**
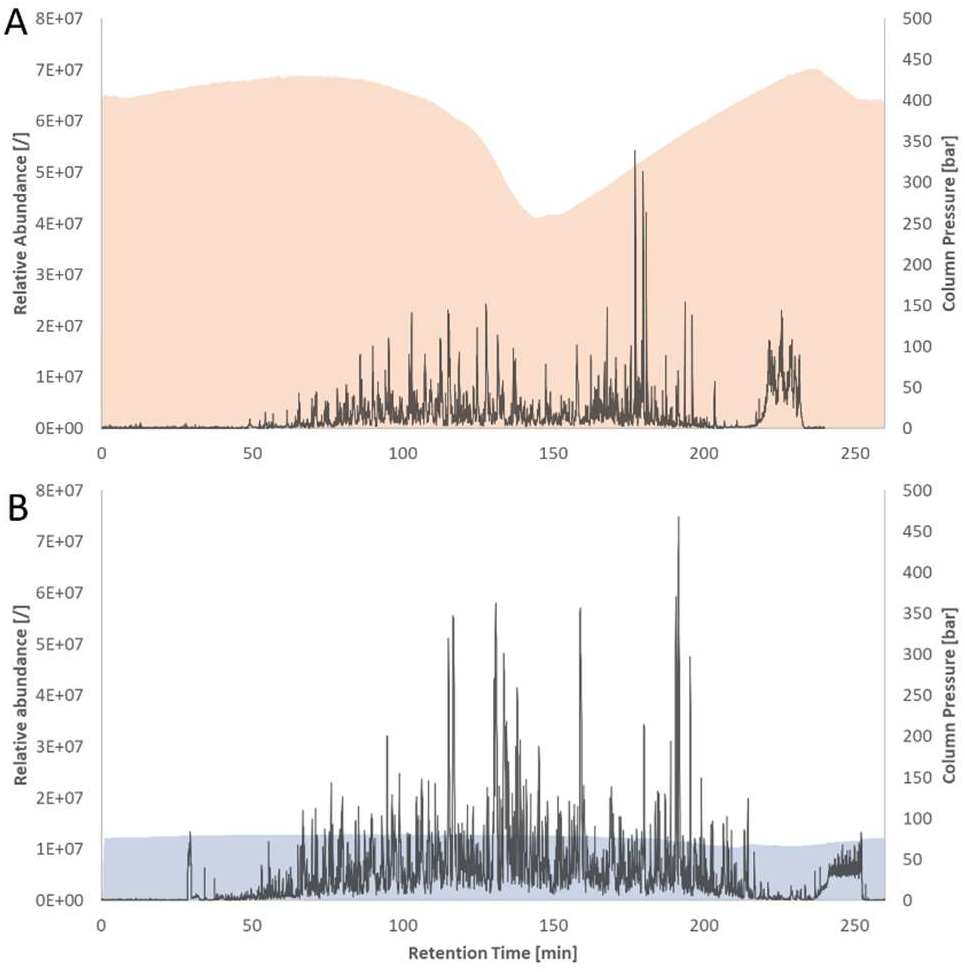
Base peak chromatograms obtained by analysis of 500 ng HEK293T cell lysate digest with a 4 hour LC-MS/MS gradient. A, (orange) obtained on a 40 cm packed bed nano LC column that was prepared in house. B, (blue) obtained on a 200 cm PharmaFluidics µPAC™ column. Corresponding column pressures are plotted on the secondary X-axis. Both columns were heated to 50°C and peptides were separated at a flow rate of 300 nl/min.

In contrast to monolithic columns, the pillar array column format has the additional advantage that perfect order is introduced into the chromatographic bed, hereby reducing peak dispersion to an absolute minimum^27^. The increased base peak intensity observed for the separation of 500 ng of HEK293T tryptic protein digest (Figure 2) already suggests that peptides have sharper elution peaks from the µPAC™ column as compared to the 40 cm packed bed nano LC column. The sharper the peptides elute from the LC column, the more concentrated these peptides enter the MS, resulting in a higher MS detector response and a better signal-to-noise ratio. In addition, sharper peptide peaks reduce peptide co-elution and result in decreased ionization suppression and hence enhanced MS sensitivity. This has a dramatic effect on the MS/MS fill times in the linear ion trap. In this experiment, a maximum time of 20 ms was allowed to achieve an AGC signal of 5e3 ions, a requirement that was easily when using the µPAC™ column, but that appeared troublesome with the 40 cm packed bed column, especially with increasing gradient length (Figure 3). Faster filling of the trap is expected to lead to more high quality MS/MS spectra with a concomitant increase in peptide and protein identifications (see below).

**Figure 3.**
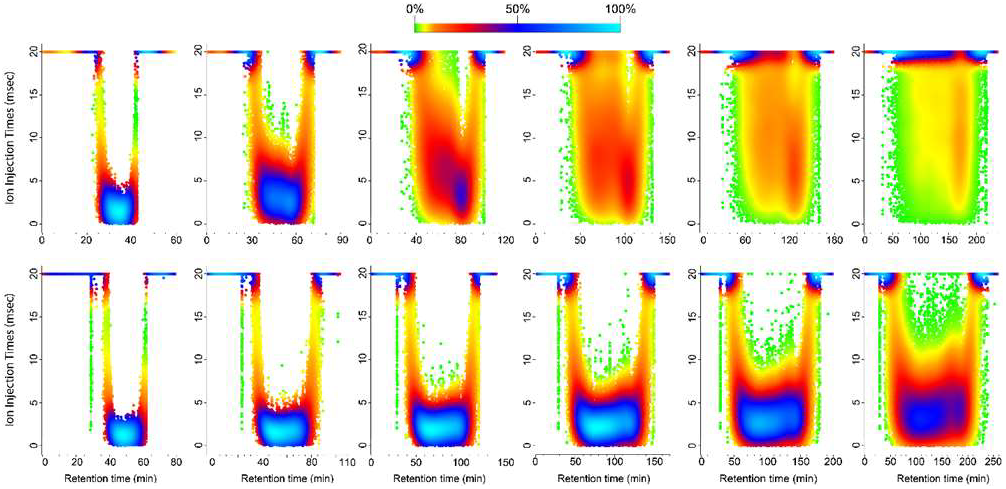
Distribution of the ion fill times in the linear ion trap for MS/MS fragmentation with a maximum fill time of 20 ms. The ion times (msec) are plotted against the retention time (min) and colored according to the density of the data points (0-100%, green-light blue). The results are shown for the 40 cm packed bed column (top) and the µPAC™ column (bottom) in combination with the 30, 60, 90, 120, 150 and 210 min gradient (left to right) for 2 µg of injected peptide material. The 40 cm packed bed column shows a rapid dispersion towards the maximum ion fill times with increasing gradient length. This effect is far more limited with the µPAC™ column where the ion times remain below 10 ms for the majority of the MS/MS scans.

The average peak width can easily be monitored throughout an entire LC-MS/MS experiment and is often used to compare the chromatographic performance of different experimental setups or column types^8^. We extracted the peak width for all identified peptides using the moFF algorithm^32^. A comparison of the peptide peak widths for both column types is shown in Figure 4A, where peptide peak widths for all sample loads are plotted as a function of the gradient time that was applied. Even though the average peak width obtained for both column types is quite similar when short solvent gradients (30 to 60 min) are used, with average peak widths ranging from 0.22 to 0.27 min, the µPAC™ column shows more narrow peaks when extending gradient times beyond 120 min. Indeed, for a 210 min gradient separation, the average peak width obtained for the packed bed column is 0.40 min compared to 0.31 min obtained on the µPAC™ column. The rate at which the peptide peak width increases with gradient time is also much higher for the packed bed column. According to the linear relationships that were found between gradient time and average peak width for both column types (*y* = 5*e*^−4^*x* + 0.2465 with R^2^ 0.87 and *y* = 3*e*^−4^*x* + 0.2482 with R^2^ 0.99 for the classical packed bed and µPAC™ respectively), this rate is almost two times higher (Figure 4B). By extending gradient times beyond 210 min, the relative difference in peak width between the two column types was expected to increase. To highlight this, three extra-long gradient durations of 330, 450 and 570 min were added. These very long gradients have only been applied for the separation of 2 µg tryptic digest on the µPAC™ column, resulting in an average peak width of 0.43 min for a 570 min gradient separation. Remarkably, a 450 min gradient separation performed on the µPAC™ column resulted in an average peak width that is comparable to what was achieved with a 210 min gradient on the 40 cm nano LC column. The improved separation performance or resolving power becomes even more apparent when peak capacities for both column types are compared (Figure 4C). Taking both the elution window and the average peak width for all identified peptides into account, a 28% increase in peak capacity for the µPAC™ column (668 versus 521) was obtained when applying a 210 min solvent gradient. Extending the gradient time to 570 min resulted in an exceptional peak capacity value of 1,331 for the µPAC™ column.

**Figure 4.**
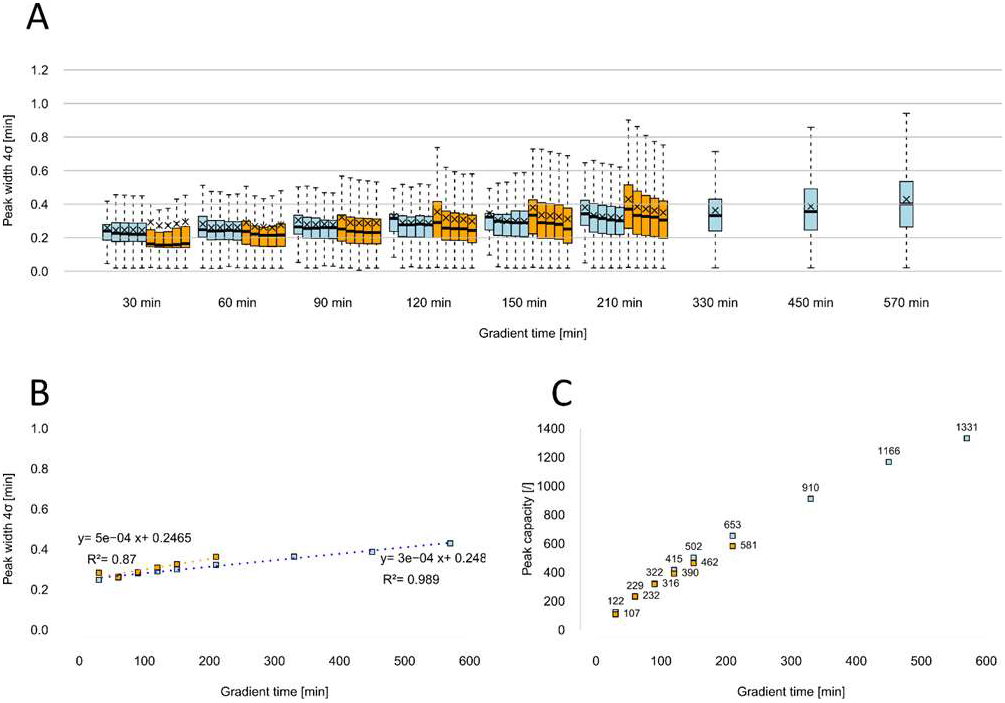
Chromatographic metrics calculated for the separation of HEK293T cell lysate digests on a µPAC™ (blue) and a packed bed nano LC column (orange). A, Peak widths (4σ, calculated from FWHM values) for all peptides that were identified and for all injected amounts. From left to right: 0.1-0.5-1-2-3 µg HEK293T digest. Average values have been marked with X. B, Average peak widths (4σ, calculated from FWHM values) plotted as a function of gradient time. Good linear relations were found for both column types. C, Peak capacity values for the separation of 2 µg HEK293T digest. Plotted as a function of gradient time.

We wondered if the increase in peak capacity translates into higher numbers of identified peptides. The average numbers of unique peptides (n=2) that were identified for the different sample loads (0.1-0.5-1-2-3 µg) and gradient durations (30-60-90-120-150-210-330-450-570 min) are given in Figure 5A. For all sample loads, consistently more peptides were identified when working with the µPAC™ column. The relative increase in peptide identifications gradually rises from 30% for short gradients to more than 50% for longer (210 min) gradients, obtaining an averaged total of 27,140 unique peptides with the µPAC™ column compared to 17,665 with the 40 cm packed bed nano LC column. The data shown in Figure 5A clearly indicate a relation between the absolute number of unique peptides that can be identified and the amount of sample that is injected. For both column types, the optimal sample load in terms of peptide identifications was reached when injecting 1 µg or more. Increasing the sample load to 2 or 3 µg did not result in an observable increase in identifications, nor did this have an drastic effect on the chromatographic performance as can be observed in Figure 4A. Even though the absolute number of peptide identifications decreases when the sample load is reduced below 1 µg, decreasing the sample load definitely has a positive effect on the relative gain that can be achieved with the µPAC™ column. At the lowest injected amount of 0.1 µg, nearly double (198%) the number of peptides could be identified using a 210 min solvent gradient, implying that the improved chromatographic separation enhances the MS sensitivity. For this sample load, increasing the gradient time beyond 210 min was not believed to yield a substantial gain in identifications, as additional dispersion would cause the concentration of low abundant peptides to drop below the limit of detection of the MS. However, even when such low amounts are used, a considerable gain in identifications can still be achieved on the µPAC™ column when increasing the gradient time. This better performance with low amounts of sample opens opportunities to analyze samples with a low protein content, such as working with a limited number of cells or a single cell samples.

**Figure 5.**
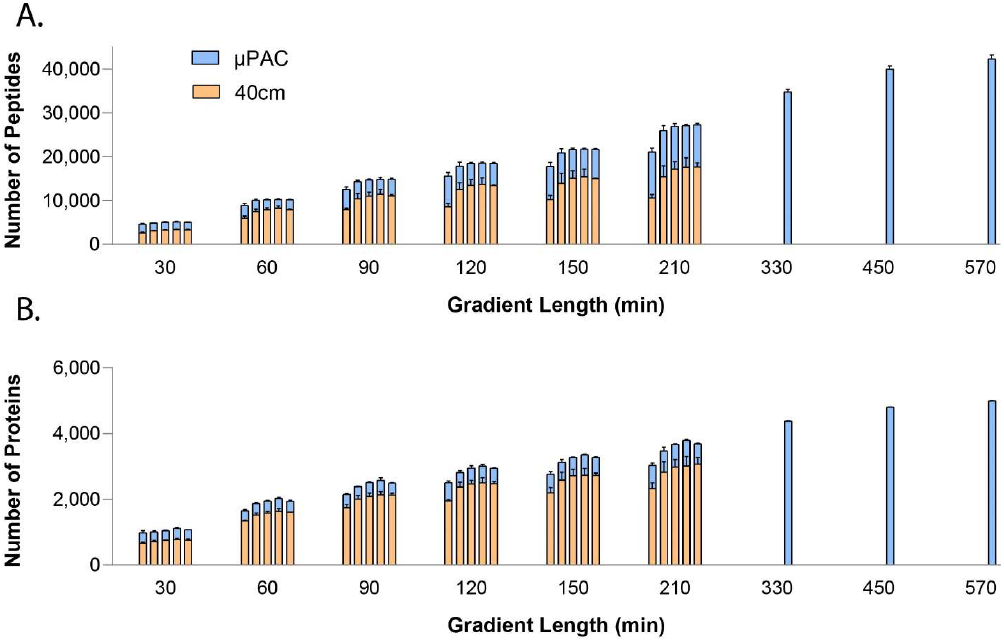
Number of identifications achieved for the separation of HEK293T digest samples on a µPAC™ (blue) and a 40 cm packed bed nano LC column (orange). A, Number of unique peptides plotted against gradient time, from left to right: 0.1-0.5-1-2-3 µg HEK293T digest. B, Number of protein groups plotted against gradient time, from left to right: 0.1-0.5-1-2-3 µg HEK293T digest.

When separating 2 µg of HEK293T tryptic digest sample using a 570 min solvent gradient, an average of 42,346 peptides were identified which corresponded to 5,004 protein groups (Figure 5B). In line with the observations on the peptide level, considerable improvements of the proteome coverage were achieved with the µPAC™ column. On average 20% more protein groups were identified compared to the packed bed nano LC column (Figure 5B). Again, the difference was higher when injecting limited sample amounts. An increase in protein group identifications up to 30% could be achieved when injecting 0.1 µg of tryptic digest.

The percentage of peptides that is shared between duplicate runs is substantially higher on the µPAC™ column, with respective peptide overlap percentages of 77% and 68% for the µPAC™ and the packed bed nano LC column (Figure S-2, Supporting Information). Combining duplicate runs, a total number of 33,501 versus 23,471 unique peptides were identified for a 210 min gradient separation. As the overlap percentage is systematically lower on the packed bed nano LC column, relatively higher profits can be made with repetitive analysis of an identical sample and combining the results on this column. In absolute numbers however, the µPAC™ column still results in a net increase in unique peptide identifications of over 40% compared to the packed bed column. Moreover, 80% of the peptides that were identified by combining packed bed nano LC results were identified when working with the µPAC™ column. In terms of protein group identifications, an even higher overlap was observed between duplicate runs, likely as additional peptides often belong to protein groups that have already been identified.

An overlap percentage of 87% was observed for the packed bed column, compared to 90% for the µPAC™ column. With a combined total of 4,048 protein groups on the µPAC™ column, this is a net increase of 16% when compared to the packed bed column. These values have been calculated for a head on comparison, where results of duplicate runs have been pooled for both columns. Comparing a single shot analysis on the µPAC™ column to the combined result of duplicates on the packed bed column, the µPAC™ column still outperforms the classical packed bed column with 15% more peptides and 6% more protein groups identified in half the analysis time. Combining the results of replicate runs is a conventional strategy to increase proteome coverage^9^, and the benefits of running several shorter gradients versus a single long gradient separation have been a topic of discussion and research in the field of proteomics^8, 34^.

The gain in identifications on a packed bed column is known to plateau from 8 h of gradient time^8^, as is shown in Figure S-3, Supporting Information. As the identification gain for the 210 min gradient remains high for the µPACTM column, we prolonged the gradient times even further. The gain in identifications was further elaborated compared to combining different LC-MS/MS runs. When combining two 210 min gradient separations, a total of 33,318 unique peptides and 4,292 protein groups were identified. By injecting the same sample and performing a single shot separation with a gradient duration of 330 min (which equals ¾ of the total analysis time) substantially more identifications were obtained. With an average of 34,834 unique peptides and 4,380 protein groups, extending the gradient time clearly is more efficient than running duplicates. These results become even more striking when the combined runs are compared to a single 450 min gradient separation. On average, 40,035 unique peptides and 4,814 protein groups could be identified in a single shot analysis. These results confirm that increasing the gradient time is a valid strategy if deeper proteome coverage is desired for the µPACTM column. However, this will only make sense if considerable gain in peak capacity is obtained by doing so. When working with the µPAC™ column, extending gradient times beyond 300 min definitely has the potential to identify unique peptides that are not likely to be identified by combining several shorter gradient runs, as a substantial gain in peak capacity can still be made in this gradient time regime.

inally, we evaluated the reproducibility of separations on the µPAC™ column by repeated injections of 25 ng of a commercial K562 cell digest supplemented with 100 fmol of 6 heavy reference peptides (QC4Life Reference Standard, Promega). These injections were carried out over a period of 6 weeks and on a single µPAC™ column connected to a Q Exactive HF mass spectrometer operated 24/7 in a core facility setting. In total, we collected data for 16 injections used as quality control runs in combination with the QCloud system. Analysis of the heavy reference peptides revealed high reproducibility in terms of retention time, full peak widths and FWHM, along with a nearly identical backpressure profile. This high level of reproducibility had a beneficial effect on label-free quantification of peptides and proteins between runs in the beginning, middle and start of the 6-week period with an average Pearson correlation of 92% and 98% for LFQ intensity values of peptides and proteins, respectively (Figure 6).

**Figure 6.**
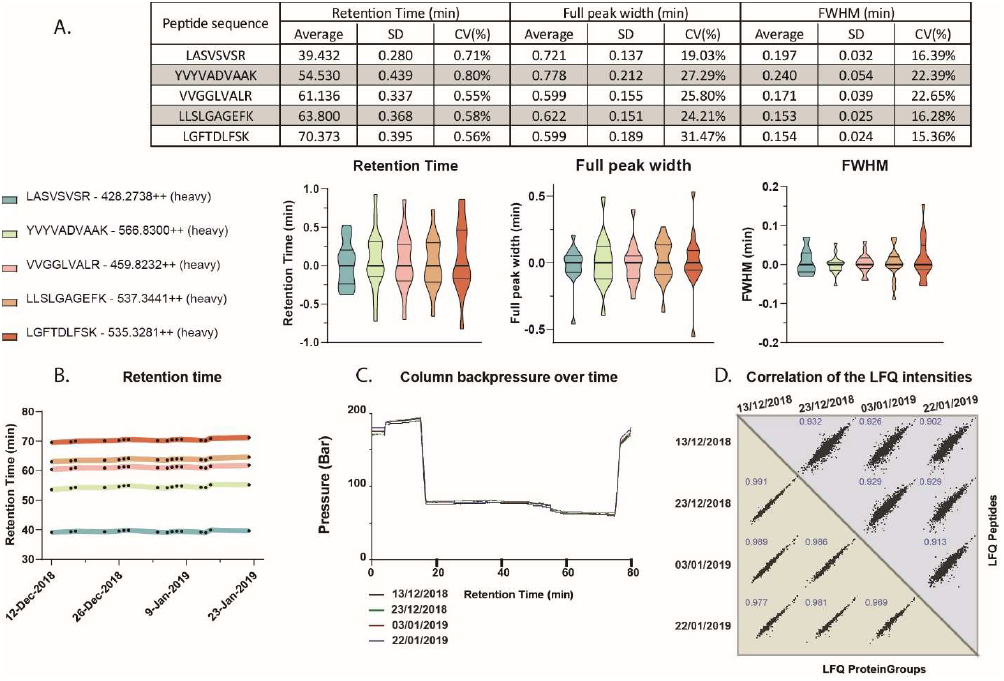
The robustness of the µPAC™ column ensures qualitative and quantitative reproducibility over time. 25 ng of a commercial K562 cell digest mixed with 100 fmol of a heavy reference peptide mix was injected 16 times over a period of 6 weeks on a µPAC™ column linked to a Q Exactive HF instrument. A, Table showing the average, standard deviation (SD) and %CV of the retention time, full peak with and full width at half maximum (FWHM) for the indicated heavy reference peptides. Violin plots representing the distribution of the normalized (median subtraction) data points (n=16) for the same parameters. B, The absolute retention times of the indicated peptides plotted over the 6-week period. C, Bbackpressure profile of four selected runs in the beginning, middle and end of the 6-week period. D, Multi-scatter plot showing the LFQ intensity values and associated Pearson correlation for quantified peptides and proteins in the four selected runs.

## Conclusions

Different approaches can be used to maximize the output of shotgun proteome profiling experiments^3–5, 34^. Compared to the strategy of performing multidimensional separation prior to MS analysis, which is associated with an increase in LC-MS/MS time and potential sample loss, our approach is substantially easier and can even be more sensitive when adequate LC-MS/MS conditions are used. As MS scanning speed is increasing progressively, the impact of the chromatographic separation quality on the depth of proteome profiling is becoming more significant. Nano LC columns are available in a wide range of formats and chemistries and are widely used among proteomics core facilities. The search for better chromatography has eventually lead to the use of relatively long columns (40-75 cm) that are packed with ≤ 2 µm silica particles. With peak capacity values reaching ~1,000, very deep proteome coverage has already been documented^9^. Further increase in chromatographic performance is however not that trivial as an increase in column length or a decrease of the packing particle diameter will have a significant impact on the LC pump pressure needed to operate these columns.

Micro pillar array columns (µPAC™) are a novel column type that shows great potential in overcoming these limitations. Micro fabrication technology allows precise control over the column geometry, enabling the fabrication of very long columns with exceptional uniformity within the separation bed or stationary phase. On top of offering unrivaled column-to-column reproducibility, very high peak capacities can be achieved at LC pump pressures below 100 bar. We investigated how these new column types improve the LC-MS/MS analysis of complex human proteomes. When comparing a 200 cm long µPAC™ column to a 40 cm packed bed nano LC column, up to 50% more peptides and 30% more protein groups could be identified in a tryptic digest of a human cell lysate. The µPAC™ column proved to yield the largest gain in identifications when limited sample amounts were injected. This increase in identifications is in line with the improved chromatographic performance of the µPAC™ column. The rate at which peak width increased according to the gradient time was found to be more than two times lower compared to a packed bed column. For a 210 min gradient, this resulted in an average peptide peak width of 0.31 min for the µPAC™ column compared to 0.40 min for the packed bed column. Extending the gradient time even further to 570 min, an average peptide peak width of 0.43 min was achieved, producing an exceptional peak capacity of 1,331 at operating pressures 6 times lower as observed on the packed bed nano LC column. An increase in run to run reproducibility is shown by a significantly higher overlap between duplicate runs for the µPACTM column. Combining several replicate runs to increase proteome coverage is a common strategy in bottom-up proteomics, however data generated in this study suggest that performing a single shot analysis on the µPAC™ column at gradient times exceeding 300 min offers deeper and more sensitive proteome coverage. Finally, the µPAC™ column showed stable performance when used in a routine setting in combination with a workhorse mass spectrometer, demonstrated by Pearson correlations of over 90% for peptide and protein intensity values in samples analyzed with 6-week time interval.

## ASSOCIATED CONTENT

### Supporting Information

Additional information as noted in text (Figures S-1-3 and Supporting Experimental Section, all as PDF files). The Supporting Information is available free of charge on the ACS Publications website.

## AUTHOR INFORMATION

### Author Contributions

The manuscript was written through contributions of all authors. All authors have given approval to the final version of the manuscript. ‡These authors contributed equally.

## ACKNOWLEDGMENT

The authors would like to acknowledge the VIB Tech Watch Fund for supporting early access to innovative technology platforms, L.M. acknowledges support by the Research Foundation − Flanders (FWO) (Project No. G.0425.18N) and F.I. acknowledges support by FWO-SBO Project S006617N.

---

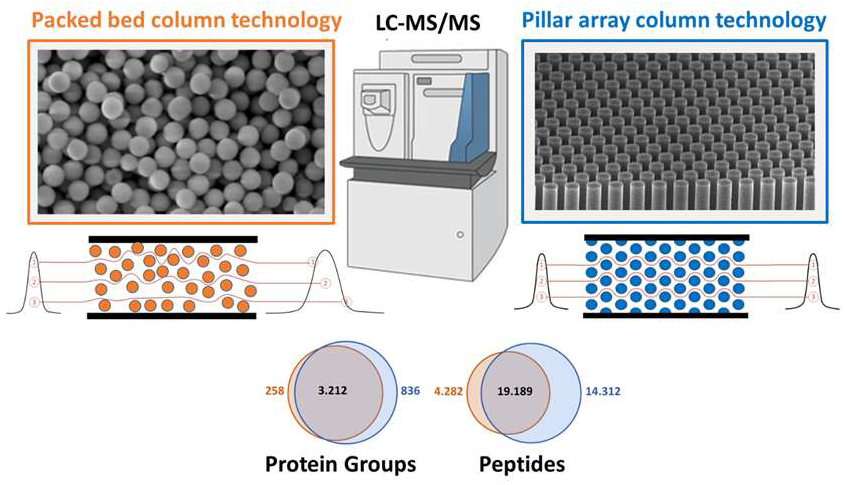

