## Supporting methods and figures for "A well-ordered nanoflow LC-MS/MS approach for proteome profiling using 200 cm long micro pillar array columns"

### Experimental Section

**Sample preparation for proteome analysis.** 10 pellets of HEK293T cells were re-suspended in 1 mL lysis buffer (8.0 M urea, 20 mM hydroxyethyl piperazineethanesulfonic acid (HEPES), pH 8.0) at room temperature (RT) and sonicated in a 10°C water bath at high intensity for 10 cycles of 30 s (Diagenode). Cell debris was precipitated by centrifugation (20,000 x g for 15 min at RT) and the supernatant was transferred to a fresh Eppendorf tube. After measuring the protein concentration (Bradford assay), cysteine residues were reduced (5 mM dithiothreitol, 30 min at 50°C) and subsequently alkylated (10 mM iodoacetamide, 15 min at room temperature in the dark). The sample was then diluted two-fold using 20 mM HEPES pH 8.0 to a urea concentration of 4 M and proteins were digested by endo-LysC (Wako, 1:100, enzyme:protein ratio) for 4 h at 37°C. The sample was further diluted using 20 mM HEPES pH 8.0 to a urea concentration of 2 M and digested with trypsin (Promega, 1:100, enzyme:protein ratio) overnight at 37°C. The next day, the digested sample was acidified with TFA to a final concentration of 1% and incubated on ice for 15 min, followed by centrifugation at 1,780 x g for 15 min at RT to remove insoluble components. Peptides were purified using SampliQ C18 columns (Agilent) with elution in 60% acetonitrile (ACN), 0.1% TFA in water. Aliquots containing approximately 500 µg of peptide material were vacuum-dried completely and stored dry at -20°C until further use. Prior to LC-MS/MS analysis, the dried peptides were re-suspended in loading solvent (2% ACN, 0.1% trifluoroacetic acid in water) and the peptide concentration for each aliquot was determined using a Lunatic instrument (Unchained Labs). Each aliquot contained around 0.55 µg/µL peptide material in a final volume of 1 mL.

**LC-MS/MS analysis.** RPLC separation of either 100 ng, 500 ng, 1 µg, 2 µg or 3 µg HEK293T peptide material was performed using a Thermo Ultimate 3000 nanoRSLC system (Thermo Fischer Scientific) coupled to an Orbitrap Elite mass spectrometer (Thermo Fischer Scientific) equipped with either one of the two column types. Depending on the mounted column, the instrument set-up was slightly altered. The first analytical column was a classical packed bed column mounted on a PneuNimbus dual-column nanoESI source (Phoenix S&T) combined with a butterfly portfolio heater (50°C, Phoenix S&T). The set-up included a 20 µL loop and all microliter pickup injections were performed using an in-house prepared trapping column (2 cm, 100 µm inner diameter) packed with ReproSil-Pur basic 5 µm silica particles (Dr. Maisch). Sample loading was performed at a flow rate of 10 µl/min during 4 min (2% ACN, 0.1% trifluoroacetic acid (TFA)) on the trapping column after which it switched in line with the analytical column for elution.

The µPAC column was mounted in the Ultimate 3000's column oven, also at 50°C. For proper ionization, a fused silica PicoTip emitter (10 µm inner diameter) (New Objective) was connected to the µPAC™ outlet union and a grounded connection was provided to this union. For this set-up we provided the system with a 1 µL loop.

Peptides were eluted from the columns using 6 different non-linear gradients of 30, 60, 90, 120, 150 or 210 min. The gradients went from 2 to 30% solvent B (80% ACN, 0.1% formic acid (FA)) in, respectively, 21, 42, 63, 84, 105 and 147 min to 56% solvent B in, respectively, 8, 16, 24, 32, 40 and 56 min, and ultimately reaching 99% solvent B in, respectively, 1, 2, 3, 4, 5 and 7 min at a flow rate of 250 nl/min. The column was then washed for 10 min at 99% solvent B, followed by re-equilibration of the system with 98% solvent A (0.1% FA in water). For the analyses on the µPAC™ column, the same gradients were applied using an increased flow rate of 300 nl/min with an additional 30 min at 98% solvent A at the end of the gradient to ensure proper re-equilibration of the column due to the larger bed volume of the µPAC™ column (total volume of 10 µl). A blank was run in between runs on the classical bed column to reduce sample carry-over, but not on the µPAC™ column as previous results showed very little to no memory effect (data not shown). An additional three extra-long non-linear gradients of 330, 450 and 570 min were performed additionally for the µPAC™ column. The gradients went from 2 to 30% solvent B in, respectively, 231, 260 and 399 min to 56% solvent B in, respectively, 88, 120 and 152 min, and ultimately reaching 99% solvent B in, respectively, 11, 15 and 19 min at a flow rate of 300 nl/min. As previously described, these runs were followed by a 10 min wash with 99% solvent B and re-equilibration of the column with 98% solvent A for 50 min.

Mass spectra were acquired on an Orbitrap Elite mass spectrometer (Thermo Fischer Scientific). The mass spectrometer was operated in data dependent, positive ionization mode, automatically switching between MS and MS/MS acquisition for the 20 most abundant peaks in a given MS spectrum. The source voltage was 3.5 kV for the classic analytical column, 2.5 kV for the µPAC column, and the capillary temperature was 275°C. In the linear trap quadrupole (LTQ)-Orbitrap Elite, full scan MS spectra were acquired in the Orbitrap (m/z 300−2,000, AGC target $3 \times 10^6$ ions, maximum ion injection time 100 ms) with a resolution of 60,000 (at 400 m/z). In parallel, the 20 most intense ions fulfilling predefined selection criteria (signal threshold of 500, exclusion of unassigned and 1 positively charged precursors, dynamic exclusion time 20 s) were then isolated in the linear ion trap and fragmented in the high pressure cell of the ion trap (AGC target $5 \times 10^3$ ions, maximum ion injection time 20 ms, spectrum data type: centroid). The CID collision energy was set to 35 V and the polydimethylcyclosiloxane background ion at 445.120028 Da was used for internal calibration (lock mass). The mass spectrometry proteomics data have been deposited to the ProteomeXchange Consortium via the PRIDE partner repository with the dataset identifier PXD011547 (Username:, password: 9ZGfEMs2).

Each of the 5 different amounts of peptide material (100 ng, 500 ng, 1 µg, 2 µg or 3 µg) was analysed in duplicate on both columns with 6 different gradients (30, 60, 90, 120, 150 and 210 min). This resulted in a total of 120 individual runs. For both columns, the sequence list was arranged in blocks which commenced with the injection of the lowest amount of peptide material using the longest gradient (i.e. 100 ng with 210 min gradient), followed by the other gradients in descending order. The analytical runs were divided in 20 blocks for database searching using the MaxQuant software as described below. Each batch contained the six runs; a certain peptide amount analysed over 6 different gradients on one type of column. Three additional searches were performed for the extra-long gradient of 2 µg peptide material.

**Database search**. All database searches were performed with MaxQuant (v1.6.1.0) using the Andromeda search engine with default search settings including a false discovery rate set at 1% on the peptide spectral matches (PSM), peptide and protein level. The mass tolerance for precursor and fragment ions was set to 20 and 4.5 ppm, respectively, during the main search. Enzyme specificity was set to trypsin, cleaving C-terminal of arginine and lysine, also allowing cleavage at Arg/Lys-Pro bonds with a maximum of two missed cleavages. The HEK293T digests were searched against the sequences of the human proteins in the Swiss-Prot/UniProtKB database from January 2018, being 20,231 protein sequences (www.uniprot.org). Variable modifications were set to oxidation of methionine and acetylation of protein N-termini. In all database searches, the matching between runs option was turned off.

**Evaluation of the durability and reproducibility of the µPAC™ column.** A commercial K562 cell digest supplemented with the 6 x 5 LC-MS/MS Peptide Reference Mix (QC4Life Reference Standard, Promega) was used as quality control (QC) to test the reproducibility of the µPAC™ column when used for routine measurements on a Q Exactive-type instrument over a 6-weeks period. For each QC run, 25 ng K562 peptides supplemented with 100 fmol heavy reference peptides was injected on an Ultimate 3000 RSLC nano LC (Thermo Fisher Scientific) in-line connected to a Q Exactive HF mass spectrometer (Thermo Fisher Scientific). The peptides were first loaded on a trapping column (made in-house, 100 µm I.D. × 20 mm, 5 µm beads) using loading solvent A (0.1% TFA in water/ACN, 98/2 (v/v)) and analyzed on a 200 cm long µPAC™ column (PharmaFluidics) with C18-endcapped functionality. Separation of the peptide material was done using a 47 min non-linear gradient going from 2-35% solvent B (0.1% formic acid in water/ACN, 20/80 (v/v)), followed by an increase to 97% solvent B in 9 min. Column temperature was kept constant at 50°C. The mass spectrometer was operated in data-dependent mode, automatically switching between MS and MS/MS acquisition for the four most abundant ion peaks per MS spectrum. Full-scan MS spectra (350-1400 m/z) were acquired at a resolution of 60,000 in the Orbitrap analyzer after accumulation to a target value of 1E6. The four most intense ions above a threshold value of 1E4 were isolated for fragmentation at a normalized collision energy of 30% after filling the trap at a target value of 1E5 for maximum 500 ms. MS/MS spectra (200-2000 m/z) were acquired at a resolution of 15,000 in the Orbitrap analyzer. The chromatographic parameters of the heavy reference peptides were analyzed using the Skyline software (v4.2.1) to extract retention time, peak width and FWHM values shown in Figure 9A and B. To identify and quantify K562 peptides and proteins, the recorded mass spectra of the 16 runs were searched together with the MaxQuant software (v1.6.2.6) using default settings, including a false discovery rate set at 1% on the PSM, peptide and protein level. The spectra were searched against all human proteins in the Uniprot/Swiss-Prot database (database release version of January 2019 containing 20,413 human protein sequences, downloaded from www.uniprot.org). The mass tolerance for precursor and fragment ions was set at 4.5 and 20 ppm, respectively, during the main search. Enzyme specificity was set as C-terminal to arginine and lysine (trypsin), also allowing cleavage at arginine/lysine-proline bonds with a maximum of two missed cleavages. Carbamidomethylation of cysteine residues was set as a fixed modification and variable modifications were set to oxidation of methionine (to sulfoxides) and acetylation of protein N-termini. Only proteins with at least one unique peptide were retained leading to the identification of 1,234 human proteins over all 16 samples. After loading the proteinGroups table from MaxQuant in the Perseus software (v1.6.2.2), only proteins identified by site, reversed database hits and potential contaminants were removed and LFQ intensities for each sample were log2 transformed to generate the scatter plots depicted in the Figure 9D. The mass spectrometry proteomics data have been deposited to the ProteomeXchange Consortium via the PRIDE partner repository with the dataset identifier PXD013235 (username:, password: FUKuDReX).

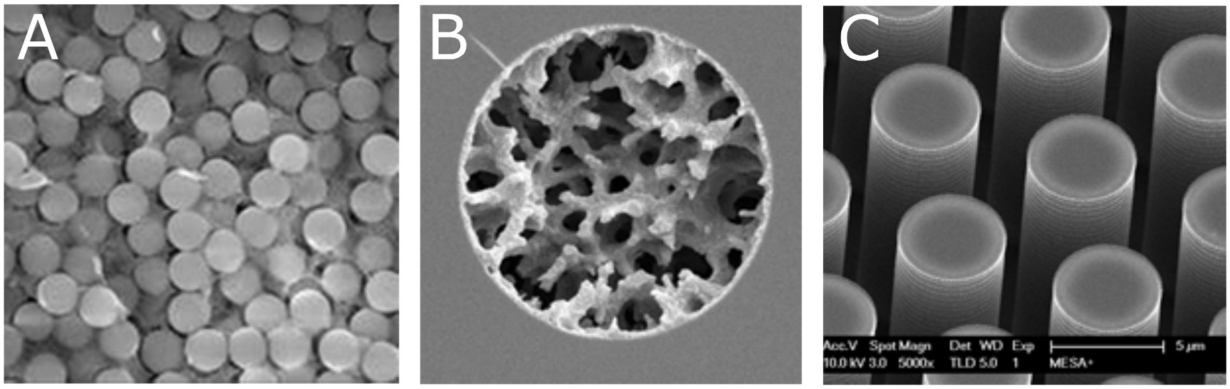


Figure S-1. Scanning electron microscopy (SEM) images showing different stationary phase support types used for the analysis of complex peptide mixtures by reversed phase liquid chromatography. A, 1.9 µm diameter silica particles, B, transverse section of a silica monolith column with an internal diameter of 25 µm, C, top view of a micro pillar array bed (5 µm diameter). Images A and B originating from G. Desmet, unpublished results.

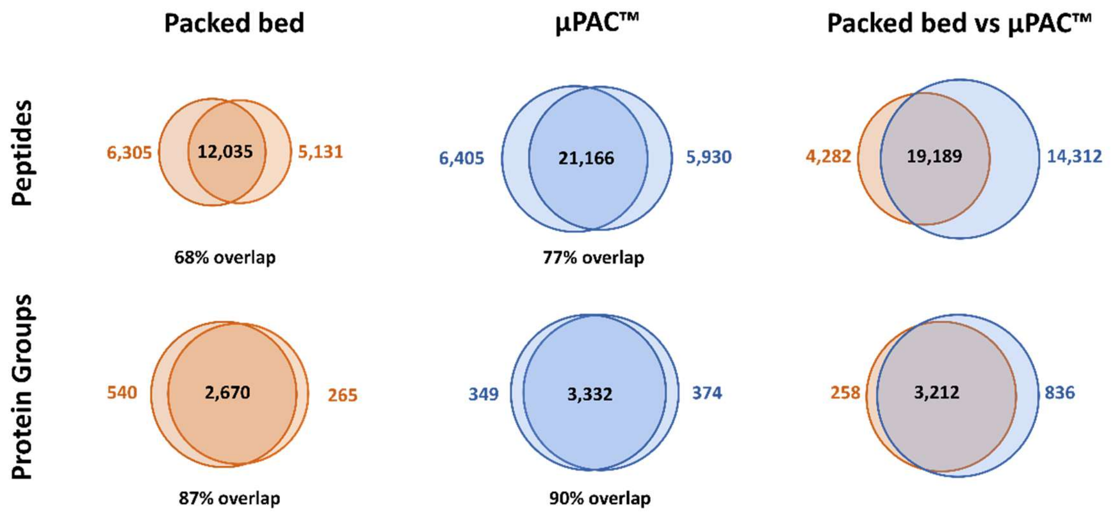


Figure S-2. Venn diagrams showing the overlap in identifications between duplicate runs and between both column types when duplicate runs are combined. 3 µg of HEK293T digest was injected on both columns and separated using a 210 min solvent gradient. µPAC™ (blue), packed bed nano LC column (orange).

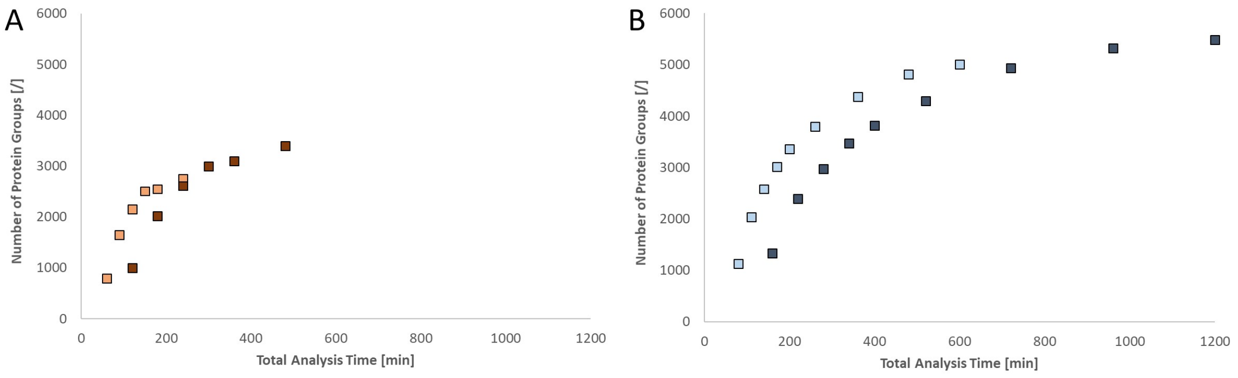


Figure S-3. Comparison of the number of protein groups that can be identified within a given time frame by running single shot analyses (light markers) and by combining the results of duplicate runs (dark markers). A, results obtained for the analysis of 2 µg HEK293T digest on a 40 cm packed bed column. B, results obtained for the analysis of 2 µg HEK293T digest on a µPAC™ column.
